# Genome-wide survey of odorant-binding proteins in the dwarf honey bee *Apis florea*

**DOI:** 10.1101/2021.02.25.432941

**Authors:** Bhavika Mam, Snehal Karpe, Ramanathan Sowdhamini

**Affiliations:** National Centre for Biological Sciences, Tata Institute for Fundamental Research, GKVK Campus, Bangalore, Karnataka, India; The University of Trans-Disciplinary Health Sciences and Technology (TDU), Bangalore, Karnataka, India

## Abstract

Odorant binding proteins (OBPs) in insects bind to volatile chemical cue and help in their binding to odorant receptors. The odor coding hypothesis states that OBPs may bind with specificity to certain volatiles and aid the insect in various behaviours. Honeybees are eusocial insects with complex behaviour that requires olfactory inputs. Here, we have identified and annotated odorant binding proteins from the genome of the dwarf honey bee, ***Apis florea*** using an exhaustive homology-based bioinformatic pipeline and analyzed the evolutionary relationships between the OBP subfamilies. Our study suggests that Minus-C subfamily may have diverged from the Classic subfamily of odorant binding proteins in insects.

## 1. Introduction

Insects are a diverse class of Arthropods with a highly sensitive olfactory system. Olfactory information helps in mate selection, oviposition while mating, foraging for food and social behaviour (Hildebrand and Shepherd, 1997). Odorant binding proteins (OBPs) abound in the sensillar lymph of insects and in the nasal mucus of many animal species with the presence of at least 50 *OBP* genes reported in some species (Hekmat-Scafe *et al*, 2002). Despite their abundance and diversity, the role of OBPs in olfactory coding is yet to be completely explored (Larter *et al*, 2016).

OBP proteins are small, soluble globular proteins, 10-30 kDa, that are further characterized by alpha-richness, and the presence of six highly conserved cysteine residues (C1-C6) with conserved disulphide spacing (Vogt et al, 1985; Pelosi and Maida, 1990) that stabilizes its tertiary structure. It has been hypothesized that OBPs bind to ligands and solubilize them to aid transport and delivery towards odorant receptors.

Genome-wide surveys to identify odorant-binding proteins in insect orders have been previously performed for various insect species in existing literature. Previous studies have predicted the presence of odorant binding proteins in various species including *Apis mellifera* (Order: Hymenoptera) (Forêt and Maleszka, 2006), *Drosophila melanogaster* (Order: Diptera) (Hekmat and Scafe, 2002; Graham and Davies, 2002), *Anopheles gambiae* (Order: Diptera) (Manoharan *et al*, 2013), *Periplaneta americana* (Order: Blattodea) (He *et al*, 2017) using homology-based bioinformatic approaches as a typical start-point.

Previous work in our laboratory (Karpe *et al*, 2016), has identified odorant receptors (ORs) in *Apis florea* using an exhaustive genomic pipeline. In order to complement the search of Ors towards a better understanding of odor coding, this study investigated odorant binding proteins (OBPs) in *Apis florea*.

*Apis florea* or the red dwarf honey bee exhibits the complex behavior of eusociality, where there is reproductive division of labour within a colony that comprises a female queen, male drones and female worker bees. While worker bees perform important tasks such as foraging, guarding the colony hive, maintenance and other diverse tasks for the colony, the queen and drone perform reproductive roles (Page and Robinson, 1991).

Members of the species exhibit haplodiploidy (Halling *et al*, 2001) system of genetic inheritance, where the males in this species are haploid, possessing half the number of chromosomes as diploid females. *Apis florea* is geographically distributed with a preference for warm climate (Otis, 1991) in regions such as mainland Asia, southern border of the Himalayas, plateau of Iran, Oman and in Vietnam, southeast China and peninsular Malaysia (Hepburn *et al*, 2005; Oldroyd and Nanork, 2009; Moritz et al, 2010) and display open nesting typically on low-lying tree branches in shaded regions (Wongsiri *et al*, 1997; Hepburn *et al*, 2005). *Apis florea* are important pollinators of tropical and ornamental plants as well as agricultural crops. They primarily feed on pollen and nectar from flowering plants. Like other honey bees, the body of *Apis florea* is studded with various types of sensilla among which olfactory sensilla (sensilla basiconica and sensilla chaetica) are prominent structures (Gupta, 1992). The antenna of the insect is typically the main site for olfactory receptors (Wigglesworth, 1965). The antennae of *Apis florea* harbor hair-like sensillae trichodea types I, II, III, IV, sensilla basiconica, sensilla placodea and sensilla ampullaceal (Gupta, 1992; Kumar *et al*, 2014).

Insect OBPs, although highly divergent, are classified on the basis of conserved cysteine signature into Classic (six cysteines), Minus-C (loss of two conserved cysteines), Plus-C (additional cysteine residues and one proline) (Zhou et al., 2004) and Atypical (∼ 10 cysteines and long C-terminus) (Hekmat-Scafe et al. 2002; Xu et al. 2003) and Dimer OBPs (two cysteine signatures). Rapid identification of repertoires of putative OBPs across various insect genomes has been suggestive of the idea that the ecological niche of an insect species may correlate with abundance of OBPs and social behaviour (Zhou et al, 2020). While reference Dipteran fruitfly, *Drosophila melanogaster* and Japanese encephalitis vector *Culex quinquefasciatus* have been found to have 51 and 110 putative OBPs respectively (Hekmat-Scafe et al., 2002; Manoharan et al, 2013), previous studies in Hymenopteran OBPs have also found species-specific differences in OBPs including 21 OBPs in eusocial *Apis mellifera* (Foret and Maleszka, 2006), 7 OBPs in fig wasp *Ceratosolen solmsi* (Wang et al, 2014) that living in closed spaces and 90 in *P. xylostella* (Vieira et al, 2012) that lives in open spaces. Using *Apis mellifera* as a closely related reference genome and a revised annotation of its OBPs, we thus investigated the identification, annotation and subfamily-based classification of putative OBPs from the genome of *Apis florea* and examined their evolutionary relationships using *in silico* approaches.

## 2. Materials and Methods

### 2.1 Obtaining genome of honeybee *Apis florea*

Aflo_1.1 genome was obtained from the National Center for Biotechnology Information (NCBI) (https://www.ncbi.nlm.nih.gov).

### 2.2 Preparing query dataset from *Apis mellifera*

AmelOBPs were pooled from the NCBI non-redundant protein database (29 putative AmelOBPs) and a previous study (Foret and Malesczka, 2007; 21 AmelOBPs) to obtain a filtered set of query protein sequences. Reciprocal homology was performed using the query set obtained and AmelOBPs from a recent study (Vieira *et al*, 2011; 21 AmelOBPs). An e-value cutoff of e^-10 was used. The resultant matches as well as unmatched OBPs (putative OBPs with no reciprocal hit; 10 protein sequences) resulted in a final dataset of annotated and unique AmelOBPs.

### 2.3 Query protein to subject genome alignments

Genomic alignments were obtained using Exonerate (Slater & Birney, 2005) with intron sizes of 500, 2000, 5000 and 10000 respectively with BLOSUM62 (Henikoff and Henikoff, 1992) as the substitution matrix.

The genomic alignments were processed as per the methodology in previous in-house study from lab (Karpe *et al*, 2016; 2017; 2020). The pipeline involves thoroughly scanning and scoring alignments to the genome based on length, degree of similarity and the best match of the scaffold location in the subject genome to the query sequence. The unique set of genomic alignments was then processed further to translate amino acids from corresponding in-frame codons. The resultant set of gene models and protein sequences were also manually corrected for missing start and stop codons, missing N-terminal and C-terminal amino acids and annotated as Complete, Partial or pseudogene.

### 2.4 Homology-based validation & nomenclature

The predicted *Apis florea* OBPs (AfloOBP) were subjected to reciprocal homology with our manually curated AmelOBP dataset, as explained above. The final dataset of predicted AfloOBPs comprised resultant matches as well as unique sequences with no corresponding hits found in the AmelOBP dataset. The AfloOBP predicted protein sequence dataset was thus annotated with respect to AmelOBP homolog, if present as well as its status as ‘**Complete**’ or ‘**Partial**’.

### 2.5 Secondary structure prediction

Secondary structure of the protein sequences were predicted using neural network-based PSIPRED v3.2 (Conesa *et al*, 2005; Buchan *et al*, 2013).

### 2.6 Signal peptide detection

N-terminal signal peptide was detected using SignalP4.1 (Nielsen et al, 1997; Petersen et al, 2011). This algorithm uses neural networks and Hidden Markov Models to determine signal peptides in a given protein sequence. The predicted signal peptide for a given sequence was cleaved off and the “mature” sequence was used for multiple sequence alignment and phylogeny.

### 2.7 Preparing dataset of insect OBPs for rooted and unrooted phylogeny

In order to prepare an outgroup for the rooted phylogeny, annotated chemosensory proteins of ***Apis mellifera*** (AmelCSPs) were obtained from a previous study (Forêt, Wanner and Maleszka, 2015), namely, AmelCSP1, AmelCSP2, AmelCSP3, AmelCSP4, AmelCSP5 and AmelCSP6.

In order to construct the phylogeny (Vogt, Große-Wilde and Zhou, 2015; Missbach, Vogel, Hansson and Große-Wilde, 2015), protein sequences of OBPs from 11 insect orders from representative insect species were obtained from previous literature and UniProt (The UniProt Consortium, 2019) database. The insect orders, corresponding species and the number of species-specific OBPs have been tabulated as in **Table 1**.

**Table 1:** List of insect orders represented in the phylogenetic tree of insect OBPs: The protein sequences of OBPs were obtained from an exhaustive literature survey referenced alongside.

| Sr.no. | Order | Reference |
| --- | --- | --- |
| 1. | Archaeognatha | Missbach *et al*, 2015 |
| 2. | Blattaria | Xu *et al*, 2009 |
| 3. | Coleoptera | Xu *et al*, 2009; Gu *et al*, 2015 |
| 4. | Diptera | Manoharan *et al*, 2013; Hekmat-Scafe *et al*, 2002,<br>NCBI |
| 5. | Hemiptera | Xu *et al*, 2009; Zhou *et al*, 2010 |
| 6. | Hymenoptera | Xu *et al*, 2009; Donnell *et al*, 2013; Gress *et al*,<br>2014; Li *et al*, 2015 |
| 7. | Isoptera | Terrapon *et al*, 2014; NCBI |
| 8. | Lepidoptera | Gong *et al*, 2009; Xu *et al*, 2009; Li *et al*, 2015 |
| 9. | Orthoptera | Xu *et al*, 2009 |
| 10. | Thysanoptera | NCBI |
| 11. | Zygentoma | Missbach *et al*, 2015 |

### 2.8 Structure-based sequence alignment and phylogenetic analysis

A structure-based seed template was obtained from the PASS2.5 database (Gandhimathi *et al*, 2012) with the SCOP ID of the fold as 47565. The dataset of “mature” AfloOBP sequences was aligned against the seed template using MAFFT (Katoh *et al* 2002; 2013). A phylogenetic tree was constructed using RaxML (Stamatakis *et al* 2006; 2014) with the maximum likelihood method with 100 bootstraps and the WAG evolutionary model (Whelan and Goldman, 2001). The phylogenetic tree was visualized and annotated using iTOL (Letunic and Bork, 2006; 2016).

## 3. Results and Discussion

We filtered and re-annotated OBPs from closely related reference genome *Apis mellifera* using a homology-based approach (See Methods). The final dataset (SI Table 1) comprised 25 AmelOBP protein sequences.

Genome-wide survey of ***Apis florea*** revealed 22 novel OBP protein sequences with 15 complete and 7 partial sequences either towards the N-terminus, C-terminus, or both with an average exon number of 5 (**Table 2; SI_Table2**). Secondary structure analysis revealed alpha-rich state of OBPs with high confidence. Typically, 6-7 alpha helices per complete AfloOBP sequence was predicted.

**Table 2:**
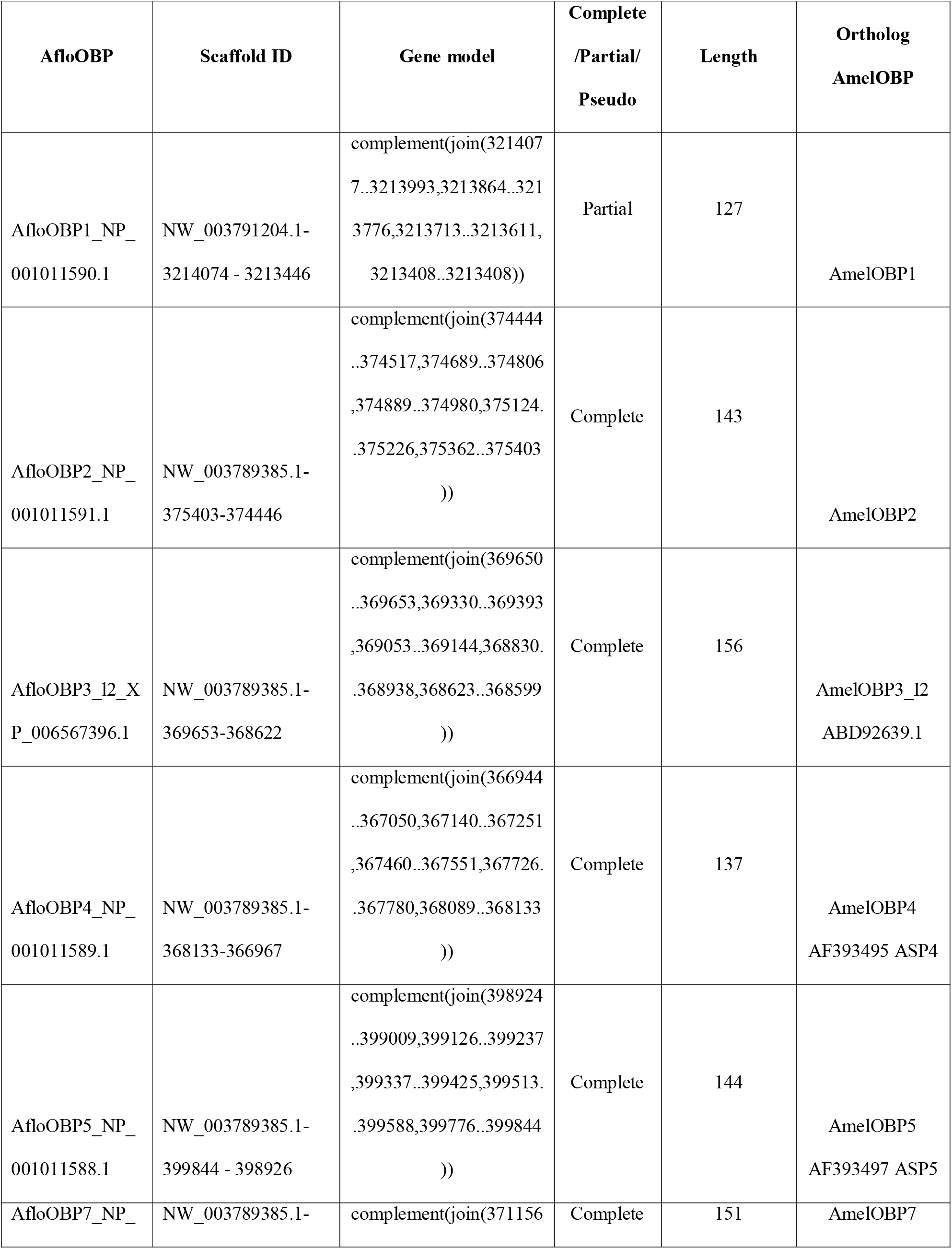
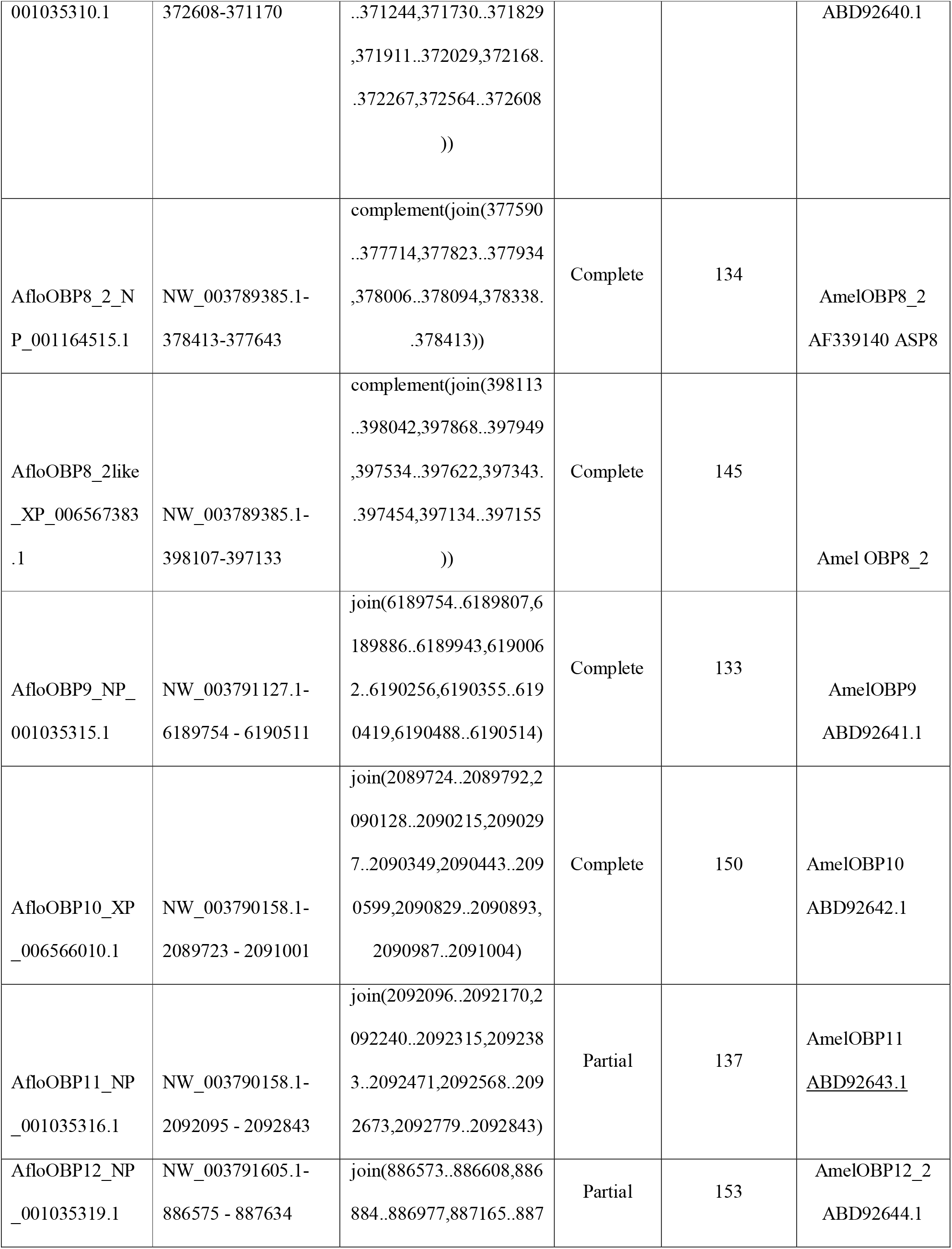
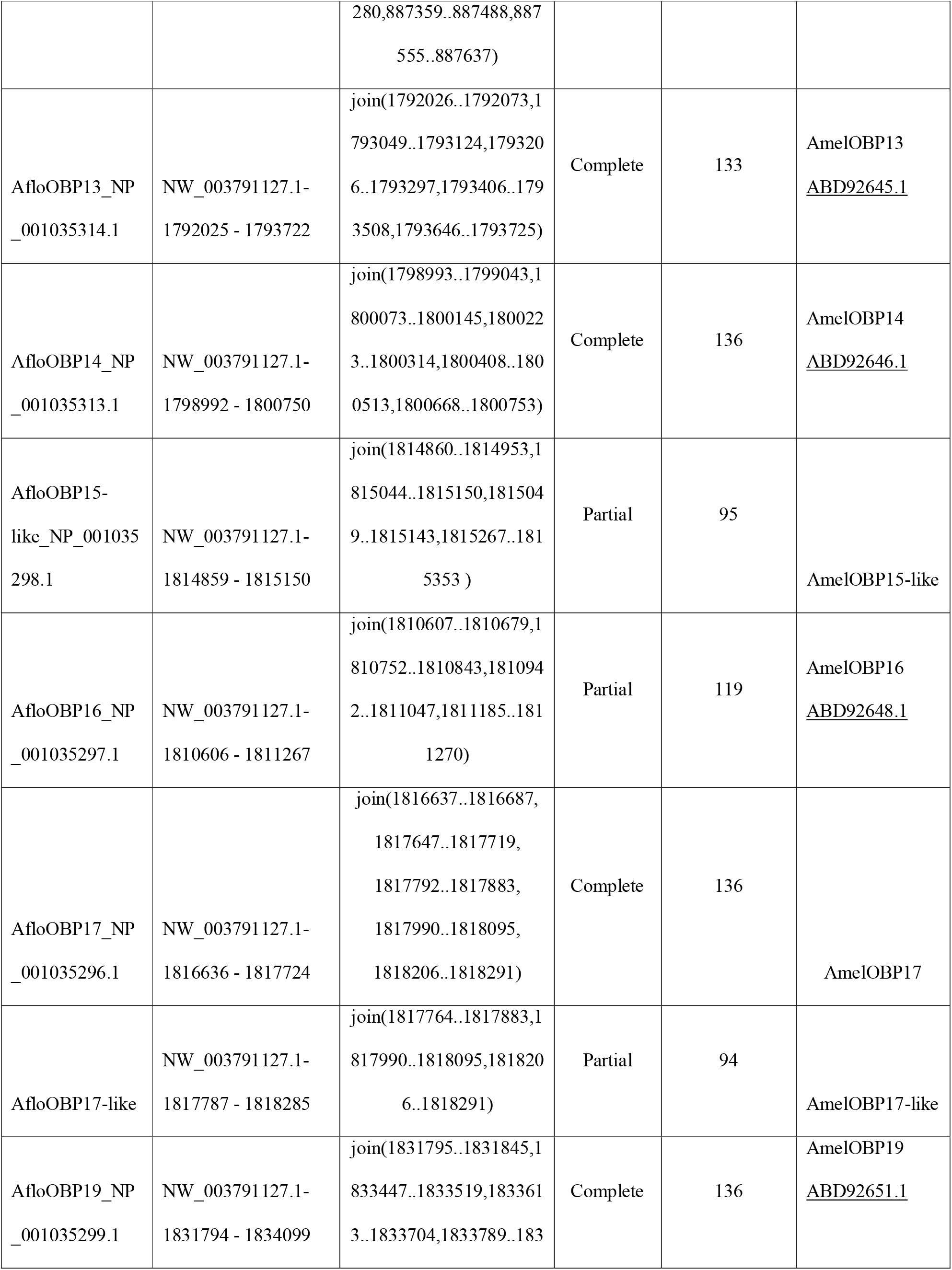
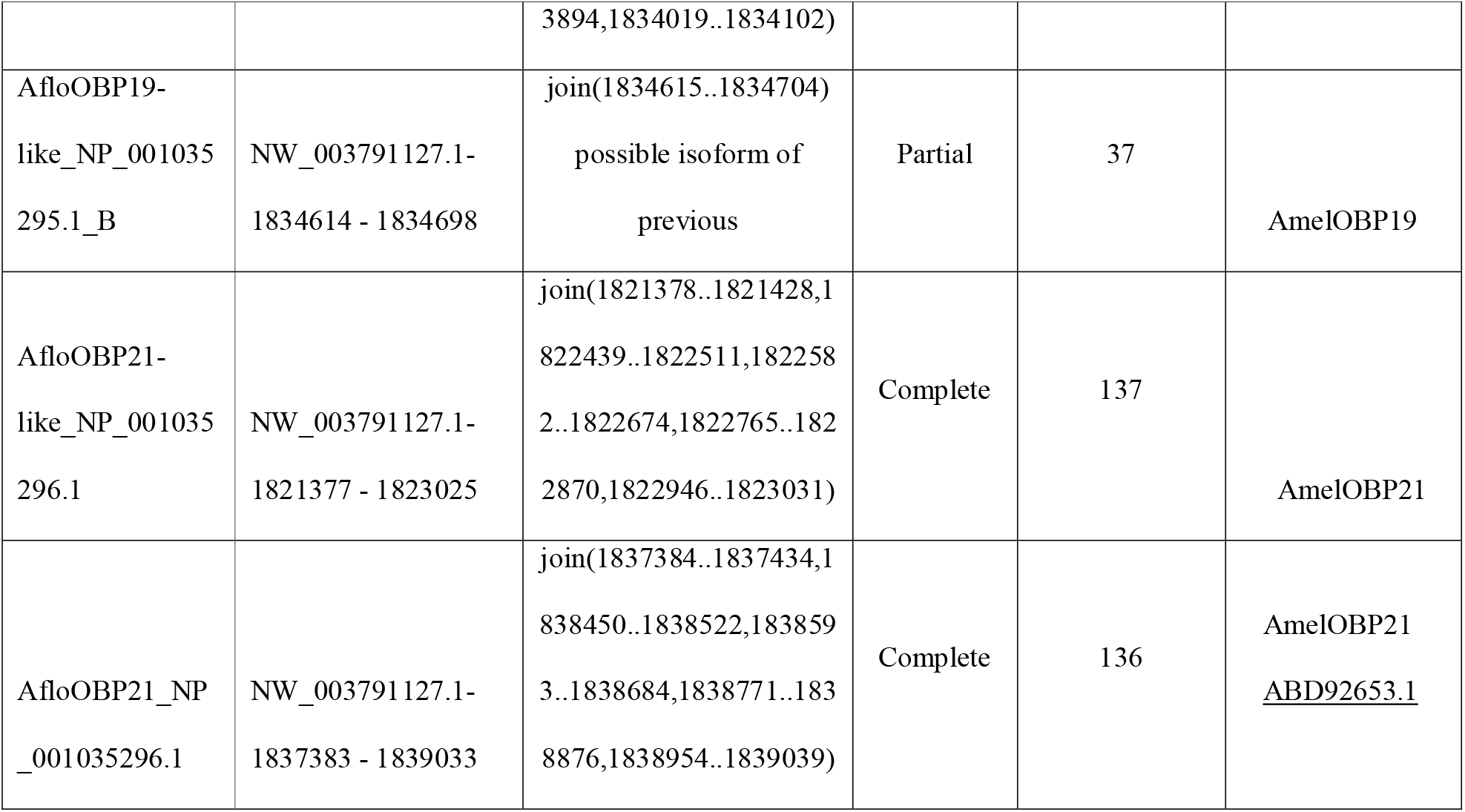
OBPs from ***Apis florea*** annotated in our study have been listed with the scaffold identity, coding exons, complete or partial status of predicted protein sequence, length of protein sequence and its ortholog in ***Apis mellifera***.

Out of 15 ***AfloOBP*** genes predicted as complete, manually corrected and annotated, 16 translated protein sequences were predicted to have signal peptide sequences. The average length of signal peptides predicted in our AfloOBP dataset was 19 amino acids. Cleavage position ranged from 16^th^ to 24^th^ amino acids in the sequence.

Sequences AfloOBP1-AfloOBP13 were found to display the conserved Cysteine signature of Classic and Minus-C subfamilies, as their orthologs in ***Apis mellifera***, and comparable to that of AgamOBP (**Figure 1)**. Multiple sequence alignment revealed conserved Cysteine profiles specific to Classic and MinusC subfamilies in the ***Apis florea*** genome (**Figure 2**). Sequences AfloOBP14-AfloOBP21 were found to show the conserved Minus-C cysteine signature where Cysteine residues in the conserved second and fifth positions are missing. Our analysis shows that the conserved cysteine signature for both subfamilies in ***Apis florea*** is similar to the representative signature observed in a previous study (Xu *et al*, 2009). The conserved cysteine signature for Classic subfamily for the Hymenopteran insect order was determined as C1-X **23:35**-C2-X**3**-C3-X 27:45-C4-X 7:14-C5-X**8**-C6 (Xu *et al*, 2009). Our study has identified 13 Classic and 9 Minus-C OBPs in ***Apis florea***. We observe the Classic cysteine signature to be conserved similarly as C1-X **27:37**-C2-X**3:4**-C3-X 33:43-C4-X 9:13-C5-X**8-9**-C6.

**Figure 1:**
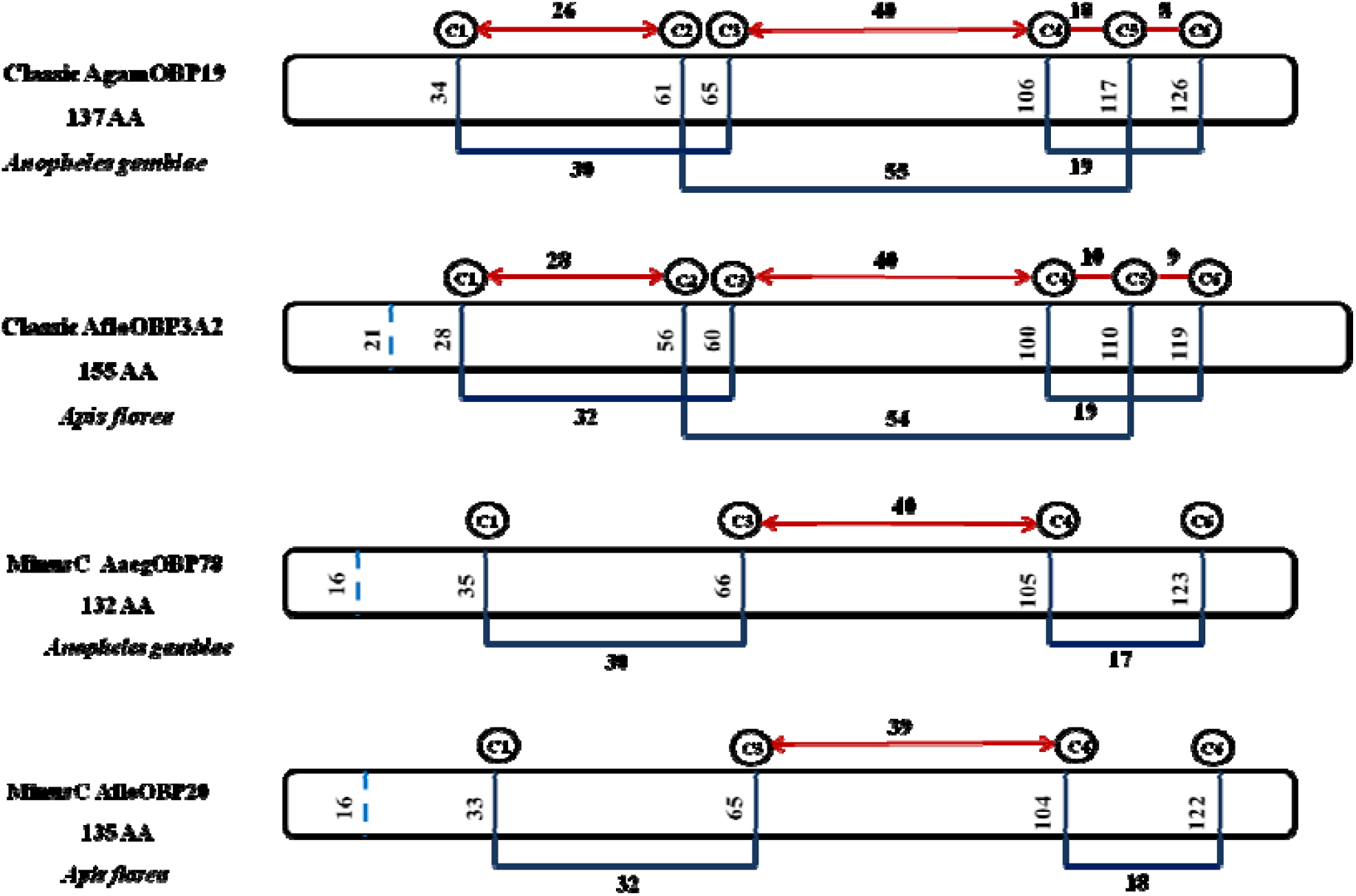
Cysteine signatures across insect orders are comparable. Cysteine residue positioning and inter-disulphide spacing is conserved in ***Apis florea*** (Hymenoptera) genome across Classic and Minus-C subfamilies. The cysteine signature of OBPs from ***Anopheles gambiae*** is given for reference.

**Figure 2:**
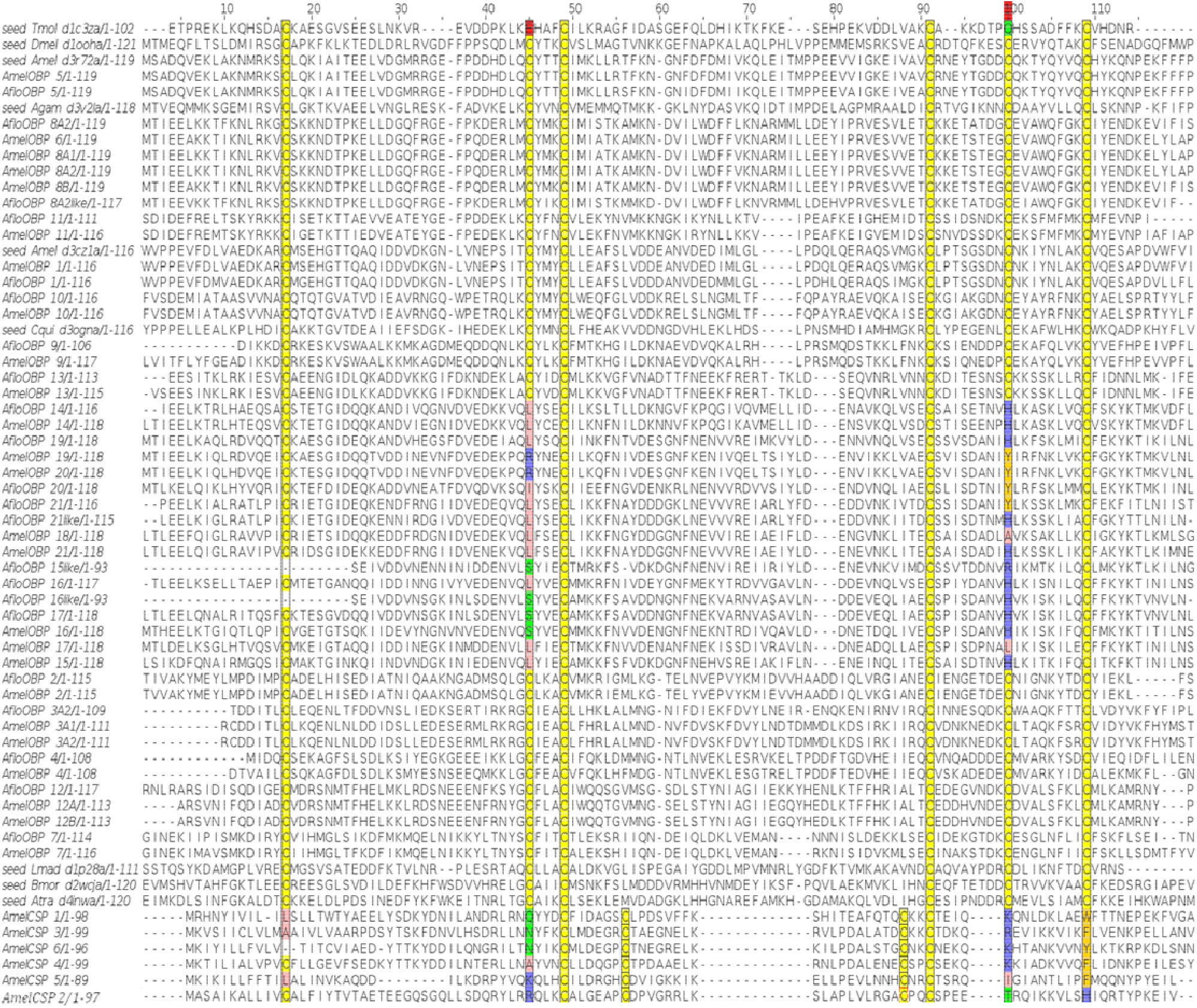
Multiple sequence alignment of OBPs from ***Apis florea***. AfloOBPs. The alignment also contains OBPs from its phylogenetic neighbour ***Apis mellifera*** (AmelOBP). Chemosensory proteins from ***Apis mellifera*** (AmelCSPs) are present as an outgroup.

Phylogenetic inference revealed the clustering of Minus-C OBPs as a sub-clade of the Classic OBP subfamily comprising members of both ***Apis mellifera*** and ***Apis florea*** OBPs (**Figure 3A**). Moreover, conserved cysteine signature specific to the chemosensory protein (CSP) family was observed in the outgroup chemosensory proteins (AmelCSP) (**Figure 2**). AmelCSPs used as outgroup clustered distinctly (**Figure 3A**) from the odorant binding proteins input to the phylogeny with 100% bootstrap value. Minus-C OBPs were found to cluster together with 60% bootstrap value closest to AfloOBP9, annotated as a Classic OBP. OBPs of the Minus-C subfamily, AfloOBP 14-20 emerge closest to AfloOBP13, a Classic OBP with an observed six cysteine signature. Interestingly, all the other Classic OBPs cluster distinctly in a clade corresponding to the insect Classic subfamily, however, AfloOBP13 clusters closely with the Minus-C group in a distinct sub-clade suggesting an evolutionary ancestral link (**Figure 3B**). Interestingly, antennal OBP (MsexABP1) from Lepidopteran insect *Manduca sexta* clustered close to the Minus-C clade along with other bee species (Hymenopteran) with high bootstrap support of 97%. It is also observed that Classic OBPs in ***Apis florea*** are phylogenetically distant from Minus-C (bee OBPs) than clades representing Atypical OBPs in Dipterans and Plus-C insect OBPs. This suggests that Minus-C OBPs in honey bees may have evolved from a single ancestral Classic OBP (similar to AfloOBP13, AmelOBP13) of its species by deletion of second and fifth cysteines. The evolution and insect order-specific occurrence of Minus-C, Plus-C and Atypical subfamilies of insect OBPs may have functional roles and would be interesting to investigate.

**Figure 3:**
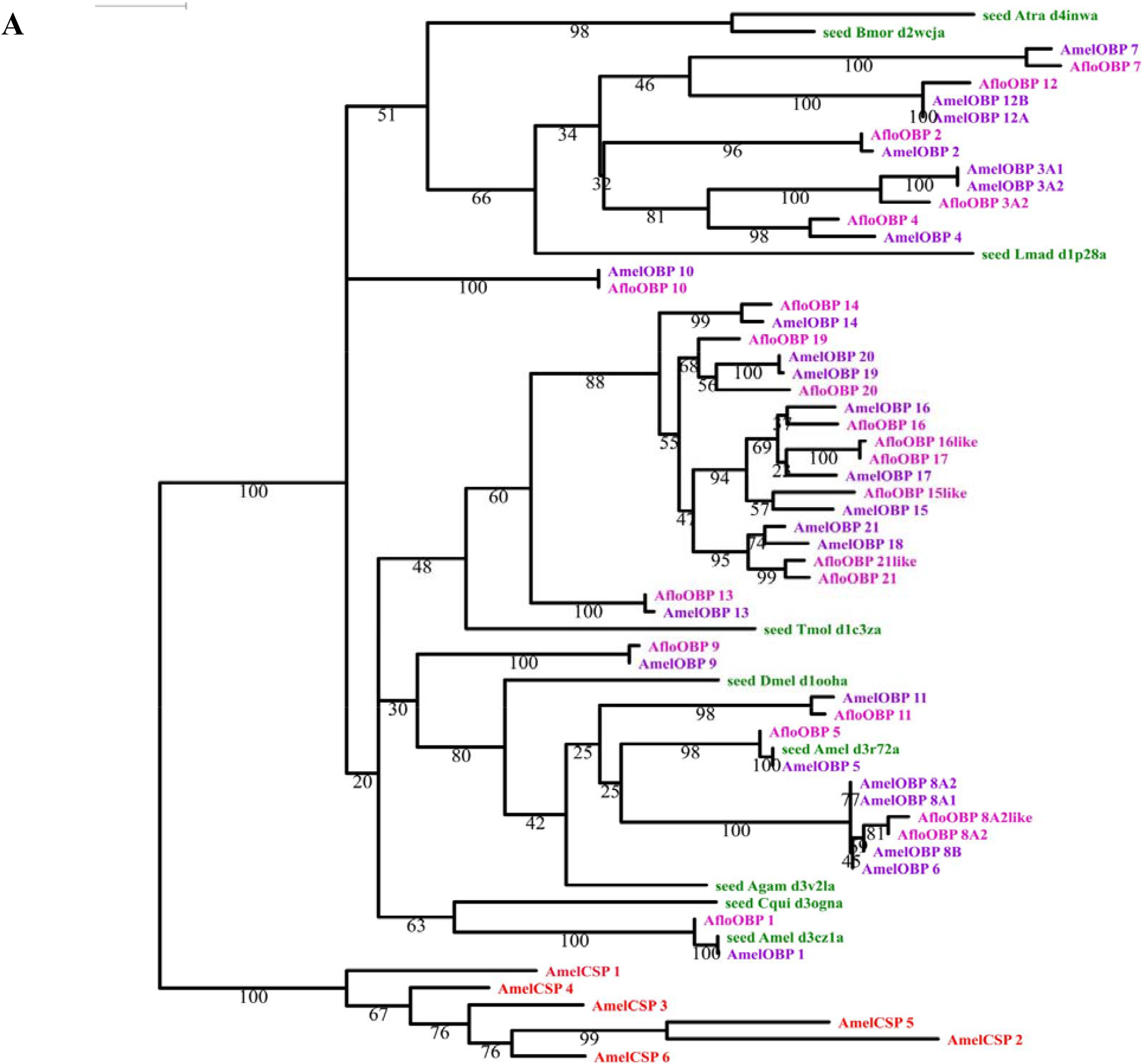
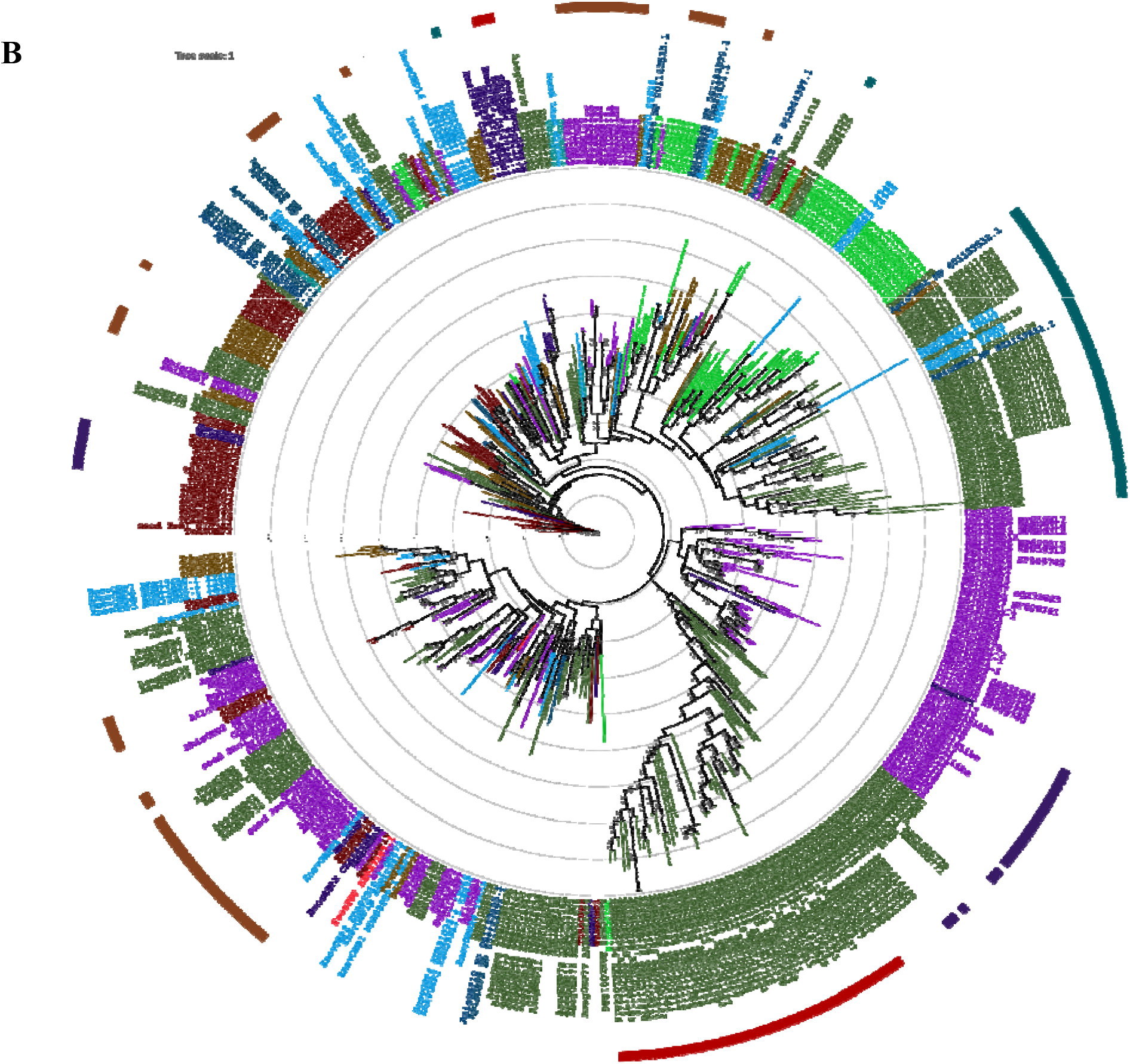
Phylogenetic tree of *Apis florea* OBPs. **Rooted phylogeny (A)** of OBPs in sister species, ***Apis florea*** (Aflo; in pink) and ***Apis mellifera*** (Amel; in purple). Members of the alignment template are colored in green, whereas the outgroup consisting of ***Apis mellifera*** chemosensory proteins (AmelCSP) is colored in red. The bootstrap values of the branches are indicated on the nodes in percentage values. **Unrooted phylogeny (B)** of OBPs from representative members of **11 insect orders** represent clade denoting Classic subfamily is colored in **brown**, Atypical in **red**, Minus-C in **violet** and Plus-C in **cyan**. The outer circle denotes members clades. The inner branch colors and label colors are colored as per order. Hymenoptera is denoted in **violet**. The bootstrap values of the branches are indicated on the nodes in percentage values.

Taken together our observations from a comprehensive bioinformatic analysis strongly suggest that Minus-C OBPs are likely to have evolved from a Classic OBP subfamily member in insects. It is possible that the evolution of a subfamily could be an adaptation to the local niche of the insect species for functional specificity (Zhou *et al*, 2020).

## Conclusion

In a step towards understanding the role of OBPs in insects, a bioinformatics-based approach was used as the starting point here. A total of 22 OBPs **including isoforms** have been identified and annotated from the genome of eusocial Asian red dwarf honeybee, *Apis florea* using a modified in-house pipeline. Our results include AfloOBPs that have been previously identified by the automated pipeline of NCBI with a query coverage and identity of 100% each (AfloOBP9 and AfloOBP11) (SI_Table3). Our annotated data includes complete OBPs that were identified as having incomplete exons in N-termini and C-termini or/and labelled as uncharacterized by the automated pipeline of NCBI. We also observe that number of *OBP* genes in *Apis florea* (22) and the western honeybee, *Apis mellifera* (25) are similar despite the differences in respective ecological niche.

We have analyzed the characteristic conserved features of these OBPs using computational methods and phylogeny resulting in discovery of new gene models as well as improvement on existing gene models from NCBI. Presence of conserved cysteine pattern, disulphide spacing, domain analysis, size and predicted secondary structure further strengthen their identity as putative insect OBPs. Moreover, the use of structurally-guided multiple sequence alignment for phylogenetic inference has been suitable

The Classic OBP subfamily clade appears to have expanded to Minus-C OBPs in honeybee and few other insect orders suggesting that Minus-C may have evolved from the Classic subfamily through strongly conserved deletions in positions corresponding to the two missing cysteines in a Minus-C OBP.

## Supplementary Materials

a. Supplementary File 1-SI_Table1 Table of OBPs from reference organism *Apis mellifera* derived from various literature sources and re-annotated to obtain a standard dataset
b. Supplementary File 2-SI_Table2 Table of predicted OBPs from *Apis florea* with annotation
c. Supplementary*_*File 3-SI_Table3 Table of blastp alignment hits obtained with query as our annotated set of AfloOBPs against Non-Redundant database.

## Ethics Approval/ declarations

Not applicable

## Data Availability Statement

All the main data generated or analysed during this study are included in this published article [and its supplementary information files]. Related datasets generated during and/or analysed during the current study are available from the corresponding author on reasonable request.

## Code availability

Not applicable

## Authors’ contributions

RS and SK conceived this research and designed experiments; BM co-designed experiments for protein sequence annotation, performed experiments, analysis, and wrote first draft of the paper. All authors read and approved the final manuscript.

## Acknowledgments

BM acknowledges Prof. (Dr.) Axel Brockmann and Prof. (Dr.) Shannon Olsson for valuable comments during the course of the study. All authors acknowledge NCBS for infrastructural support.

